# Lower motor performance is linked with poor sleep quality, depressive symptoms, and grey matter volume alterations

**DOI:** 10.1101/2024.06.07.597666

**Authors:** Vincent Küppers, Hanwen Bi, Eliana Nicolaisen-Sobesky, Felix Hoffstaedter, B.T. Thomas Yeo, Alexander Drzezga, Simon B. Eickhoff, Masoud Tahmasian

## Abstract

Motor performance (MP) is essential for functional independence and well-being, particularly in later life. However, the relationship between behavioural aspects such as sleep quality and depressive symptoms, which contribute to MP, and the underlying structural brain substrates of their interplay remains unclear. This study used three population-based cohorts of younger and older adults (n=1,950) from the Human Connectome Project-Young Adult (HCP-YA), HCP-Aging (HCP-A), and enhanced Nathan Kline Institute-Rockland sample (eNKI-RS). Several canonical correlation analyses were computed within a machine learning framework to assess the associations between each of the three domains (sleep quality, depressive symptoms, grey matter volume (GMV)) and MP. The HCP-YA analyses showed progressively stronger associations between MP and each domain: depressive symptoms (unexpectedly positive, r=0.13, SD=0.06), sleep quality (r=0.17, SD=0.05), and GMV (r=0.19, SD=0.06). Combining sleep and depressive symptoms significantly improved the canonical correlations (r=0.25, SD=0.05), while the addition of GMV exhibited no further increase (r=0.23, SD=0.06). In young adults, better sleep quality, mild depressive symptoms, and GMV of several brain regions were associated with better MP. This was conceptually replicated in young adults from the eNKI-RS cohort. In HCP-Aging, better sleep quality, fewer depressive symptoms, and increased GMV were associated with MP. Robust multivariate associations were observed between sleep quality, depressive symptoms and GMV with MP, as well as age-related variations in these factors. Future studies should further explore these associations and consider interventions targeting sleep and mental health to test the potential effects on MP across the lifespan.

## Introduction

Motor performance (MP), including dexterity, strength, endurance, and processing speed, is a fundamental aspect of human functioning. Impairment of MP predicts functional dependency in older adults (D. X. M. Wang et al., 2020), reduced quality of life and well-being of patients with neurological and psychiatric disorders (Chapuis et al., 2005; Ramari et al., 2020; Skinner et al., 2015), and overall mortality (White et al., 2013). Identifying the factors contributing to MP and elucidating the neurobiological substrates of their interplay in the general population and clinical samples has become of great interest.

Previous studies demonstrated the role of various demographic, physical, psychosocial, and lifestyle aspects on MP (Lin et al., 2020; Sternäng et al., 2015; Zeiher et al., 2019). Among them, poor sleep and depressive phenotype are highly prevalent nowadays (Chunnan et al., 2022; Joo, 2022) and strongly interrelated (Cheng et al., 2018; Olfati et al., 2024). These aspects adversely affect physical and brain health (Tahmasian et al., 2020; Y. Wang et al., 2024), and have been suggested as potential contributors to abnormal MP (T. Y. Wang et al., 2018). For example, acute sleep loss affects the MP of athletes, leading to reduced performance in the afternoon of the following day (Craven et al., 2022). Several pieces of evidence point to potential links between muscle strength and sleep duration (H.-C. Chen et al., 2017; T. Y. Wang et al., 2018), although the results remained inconclusive in a meta-analysis (Pana et al., 2021). In addition, decreased depressive symptoms is associated with higher MP, such as grip strength and cardiovascular fitness in the general adult population (Gu et al., 2021; Kandola et al., 2020; Sui et al., 2009). Thus, it is important to assess the combined role of sleep disturbance and depressive symptoms on MP, which are common stressors in the general population and co-occur in chronic clinical conditions involving motor dysfunction, including neurodevelopmental disorders, major depressive disorder, Parkinson’s disease, Alzheimer’s disease, and stroke (Lieberman, 2006; Moran et al., 2005; Shura et al., 2017; Suh et al., 2014). Hence, given the relevance of the relationship between depressive symptoms and sleep disturbance with a range of disorders and quality of life, understanding their interplay with MP will guide the development of novel preventive strategies and therapeutic approaches.

Neuroimaging studies identified brain structure correlates linking larger grey matter volume (GMV) with enhanced MP across various cortical, subcortical, and cerebellar regions (Koppelmans et al., 2015; Seidler et al., 2010). Increased GMV in frontal and parietal areas has been observed among physically active individuals (Eyme et al., 2019). Furthermore, grip strength has been associated with GMV variations in subcortical, limbic, and temporal regions (Jiang et al., 2022). However, the relationship between sleep quality and depressive symptoms with regional GMV remains inconclusive. Large-scale studies identified a link between longer sleep duration and higher GMV in basal ganglia but have failed to establish significant associations between other sleep health dimensions and GMV, such as insomnia (Schiel et al., 2023; Weihs et al., 2023). Similarly, investigations on depressive symptoms revealed either minimal effects or no significant associations between depressive symptoms/clinical depression and brain structure (for the ENIGMA-Major Depressive Disorder Working Group et al., 2016, 2017; Olfati et al., 2024; Winter et al., 2022, 2024). Taken together, the neurobiological correlates of the interplay between sleep quality and depressive symptoms with MP remain unclear. Hence, this study aims to investigate the multivariate associations across sleep quality, depressive symptoms, and brain structure with MP.

Ageing shows notable effects not only on MP but also on sleep, depressive symptoms, and brain structure. Age-related changes can be observed for motor function, such as slower movement speed and coordination (Jiménez-Jiménez et al., 2011; Seidler et al., 2010), a more fragmented sleep pattern and worse sleep quality (Mander et al., 2017), a change in depressive symptomatology and related factors, such as loneliness (Hegeman et al., 2012; Lee et al., 2021), and atrophy in regional and global GMV (Bethlehem et al., 2022; Good et al., 2001). Considering these omnipresent age-related variations within the mentioned domains, we conducted separate analyses for young and older adults to assess the differential impact of ageing on the multivariate association between them. We hypothesised that 1) stronger MP is associated with better sleep quality and less depressive symptoms; 2) this relationship is anchored in measurable parameters of macroscale brain structure; 3) a combination of sleep quality, depressive symptoms, and brain volumes has the strongest association with MP; and 4) such multivariate associations are different between young and older adults. Data from three independent publicly available cohort studies, the Human Connectome Project Young Adult (HCP-YA), the Human Connectome Project Aging (HCP-A), and the enhanced Nathan Kline Institute-Rockland Sample (eNKI-RS), were used to test the replicability of our findings. We calculated regularised canonical correlation analyses (rCCA) to explore the link between individual domains (i.e., sleep, depressive symptoms, GMV), and their combinations in association with various MP domains (i.e., dexterity, strength, endurance, and processing speed).

## Methods

### Participants and phenotypic data

We used data from 1,950 individuals across three openly available imaging cohorts and split them into four samples of younger adults (HCP Young: 22-37 years, n=1,086, 587 females; eNKI-RS Young: 18-40 years, n=230, 128 females) and older adults (HCP Aging: 50-85 years, n=354, 198 females; eNKI-RS Old: 50-85 years, n=280, 199 females) (Bookheimer et al., 2019; Nooner et al., 2012; Van Essen et al., 2013). Across all four samples, sleep quality was measured with the Pittsburgh Sleep Quality Index (PSQI) (Buysse et al., 1989), and depressive symptoms were measured with items of the Adult Self Report (ASR) associated with depression (Achenbach & Rescorla, 2003). MP was assessed through different motor tasks that varied across the cohorts. The HCP Young and Aging cohorts used NIH toolbox assessments of Grip Strength, the 2-Minute Walk Test for endurance, Pattern Comparison Processing Speed Test, with an additional 9-Hole Pegboard Test for dexterity in HCP Young and Trail Making Test A for sensorimotor speed in HCP Aging (Reitan, 1992; Reuben et al., 2013; Weintraub et al., 2013). The eNKI-RS Young and Old assessment included grip strength, VO2max by bike test for endurance, Grooved Pegboard Test for dexterity, Penn Computerized Neurobehavioral Battery (CNB) - Mouse Practice task for processing speed, and Trail Making Test A for sensorimotor speed (Åstrand & Ryhming, 1954; Delis et al., 2001; Gur, 2001; Reuben et al., 2013) (see Supplementary file for more details).

We combined HCP Young and HCP Aging to identify potential age group-independent effects based on the common motor tasks. Combining all three cohorts was impossible due to the different motor tasks across the HCP and eNKI-RS cohorts.

### MRI preprocessing & GMV analysis

T1 weighted scans of all cohorts were pre-processed using the Computational Anatomy Toolbox Version 12.8.2 (CAT12) (Gaser et al., 2022). T1 weighted scans were acquired using a MPRAGE sequence in all three cohorts. In HCP-YA this was done using a single Siemens 3T Skyra scanner with a resolution of 0.7mm isotropic voxels (Van Essen et al., 2013). In the HCP Aging cohort data was acquired across four sites using Siemens Prisma 3T MRI scanners with a resolution of 0.8mm isotropic voxels (Bookheimer et al., 2019). In the eNKI-RS data was acquired using Siemens 3T Tim Trio scanner with a resolution of 1mm isotropic voxels (Nooner et al., 2012). The mean GMV for each participant was estimated in native space, as implemented in the region of interest analysis by CAT12. To cover the whole brain, we used a combination of three atlases. The Schaefer atlas for 200 cortical areas, the Melbourne subcortex atlas scale II for 32 subcortical regions, and the spatially unbiased atlas template of the cerebellum and brainstem (SUIT) for cerebellar regions, resulting in a total of 262 regions (Diedrichsen et al., 2009, 2011; Schaefer et al., 2018; Tian et al., 2020) The images underwent an automatic quality control by CAT12; all scans with an image quality rating (IQR) of > 3.5 were excluded.

### Regularized Canonical Correlation Analysis (rCCA)

To uncover the multivariate relationship between sleep quality, depressive symptoms, and brain structure with MP, we computed five different regularised canonical correlation analyses (rCCA) for each sample in the main analysis, as summarised in Figure 1. Canonical correlation analysis assesses the links between two sets of variables by identifying the linear combinations of variables within each set, such that the resulting linear combinations are maximally correlated between sets. In this context, a *canonical variate* refers to a specific linear combination, and a *mode* represents a pair of these linear combinations, one from each variable set, that are maximally correlated. We used a regularised version with L2 penalty function to address the high dimensionality and potential multicollinearity among the large number of variables, particularly from the GMV variables. The L2 regularisation stabilises the results by penalising the magnitude of the coefficients, thereby reducing the risk of overfitting and the impact of multicollinearity (Mihalik et al., 2022; Vinod, 1976). To control for potential confounding factors, we regressed out age, age squared, and sex in all rCCA models in a cross-validation consistent manner avoiding data leakage (Sasse et al., 2024) . Additionally, to ensure that the findings were not biased by variation in head size, the total intracranial volume was regressed out for the rCCA models that included GMV. The cohort was included as a confounding factor in the supplementary analysis of the combined HCP Young and HCP Aging datasets.

**Figure 1.**
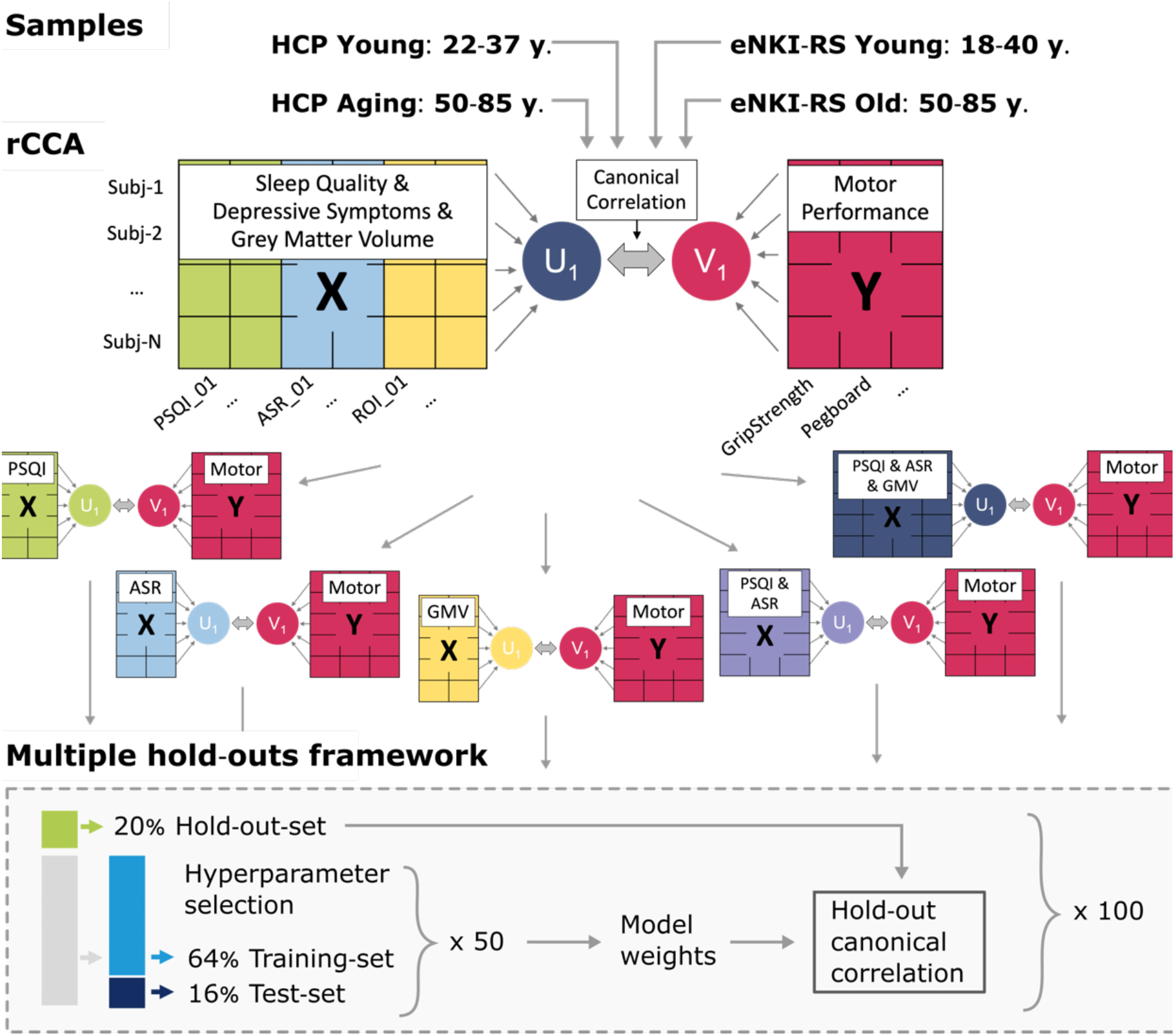
Flowchart of the analysis pipeline. Three cohorts were split into two samples of young (18-40 years) and two samples of older (50-85 years) participants. For each sample, five different regularized canonical correlation analyses (rCCA) for individual and combined domains were computed and tested in a machine learning framework. HCP, Human Connectome Project; eNKI-RS, enhanced Nathan Kline Institute Rockland Sample; PSQI, Pittsburgh Sleep Quality Index; ASR, Items of the Adult Self Report questionnaire associated with depressive disorder; GMV, Grey matter volume.

In the present study, domain X included either (1) depressive symptoms, (2) sleep quality, (3) brain structure, (4) a combination of depressive symptoms and sleep quality, and (5) a combination of depressive symptoms, sleep quality, and GMV. Domain Y included MP. The pairs of canonical variates describing the relationship between each domain X with domain Y are computed and tested for stability and generalisability using a machine learning framework that uses multiple holdouts of the data (see following section) (Figure 1). Of note, we only presented the first mode of association, as all other modes showed low generalisation to the hold-out data (data not shown). To interpret the calculated modes of the rCCA models, we calculated *canonical loadings*. These are Pearson correlations of the input variables with the canonical variates calculated in rCCA and can be interpreted as factor loadings.

### Robustness and generalisability of rCCA models

To assess the robustness of our rCCA models, we applied a multiple hold-out framework (Mihalik et al., 2022; Monteiro et al., 2016). Specifically, we divided the data into a 20% hold-out set and 80% optimisation set (referred to as the outer split). The optimisation set was then further divided into 80% for training and 20% for testing (referred to as the inner split) (Nicolaisen-Sobesky et al., 2022). The optimal rCCA hyperparameters were estimated by computing the Euclidean distance in a two-dimensional space of the stability, the similarity of weights, and the canonical correlation coefficients, of the measured vs. the perfect stability and correlation (Mihalik et al., 2020). This was done in the inner split and repeated 50 times. Subsequently, the hold-out set was projected onto the weights of the optimal rCCA model obtained from the inner split to estimate the hold-out canonical correlation. To ensure the robustness of the found associations, the outer split was randomly repeated 100 times. The hold-out correlations and canonical loadings reported in our analyses represent averages across all 100 repeats. To take into account the family structure within HCP Young, the members of a family were kept in the same data split, as implemented using the exchangeability blocks within the CCA/PLS toolkit (Mihalik et al., 2022; Winkler et al., 2015).

### Comparison of hold-out canonical correlations between rCCA models

To facilitate comparisons between rCCA models within a sample across different domains, hold-out correlations were compared. We Fisher z-transformed the canonical correlation coefficients of the first mode and computed pairwise comparisons between all domains. Given the interdependence between the optimisation and hold-out sets across repetitions, variance may be overestimated (Nadeau & Bengio, 1999). Therefore, we adjusted the t-tests according to the approach described by Bouckaert & Frank, 2004. p-values for all comparisons within one rCCA model are corrected using False Discovery Rate (FDR). P < 0.05 is considered as a significant difference between rCCA models.

Redundancy indices for the first mode of Y were computed as an additional metric for model comparison. The redundancy index quantifies how much variance in the variables set Y can be explained by the canonical variate derived from X. This is done by computing the canonical loadings for Y, squaring these loadings, and averaging them. This amount of shared variance is multiplied by the coefficient of determination from the canonical correlation to obtain the redundancy index.

### Cross-cohorts replicability

To address the difficulties replicating results in the field of cognitive neuroscience, we performed cross-cohort comparisons to ensure the replicability of our findings through qualitative replication (Xu et al., 2024). The models calculated in this study use the same questionnaires for sleep quality and depressive symptoms, as well as the same calculations for GMV. To compare the rCCA models, Spearman correlations were computed between the averaged loadings of (1) GMV and (2) PSQI and ASR separately. However, as the measures of MP differed, the comparisons remained conceptual. The first mode of each sample was compared with one another, and p-values were corrected for multiple comparisons using FDR correction.

### Ethics

The ethics protocols for analysing data from the different cohorts were approved by the Ethics Committee of Heinrich Heine University Düsseldorf (Approval No. 4039). All participants involved in the original studies, HCP, and eNKI-RS, provided informed consent.

## Results

Consistent and replicable multivariate associations of depressive symptoms, sleep quality, and GMV with MP were observed in the HCP Young, eNKI-RS Young, and HCP Aging samples. Conversely, the eNKI-RS Old sample failed to generalise to the hold-out set in all rCCA models, reflecting more variability according to our results (Figure 2a-d). Relatively modest but positive associations were observed between depressive symptoms and MP, with Fisher Z transformed hold-out correlation coefficients of 0.13 ± 0.06 (HCP Young), 0.12 ± 0.12 (eNKI-RS Young), 0.13 ± 0.1 (HCP Aging), which was unexpected given the typically reported negative impact of depressive symptoms on motor performance. Sleep quality showed slightly stronger associations with MP, yielding Fisher Z transformed hold-out correlation coefficients of 0.17 ± 0.05 (HCP Young), 0.18 ± 0.14 (eNKI-RS Young), 0.14 ± 0.1 (HCP Aging). GMV consistently showed robust associations with MP across three samples, with Fisher Z transformed hold-out correlation coefficients of 0.19 ± 0.06 (HCP Young), 0.27 ± 0.11 (eNKI-RS Young), 0.26 ± 0.11 (HCP Aging).

**Figure 2.**
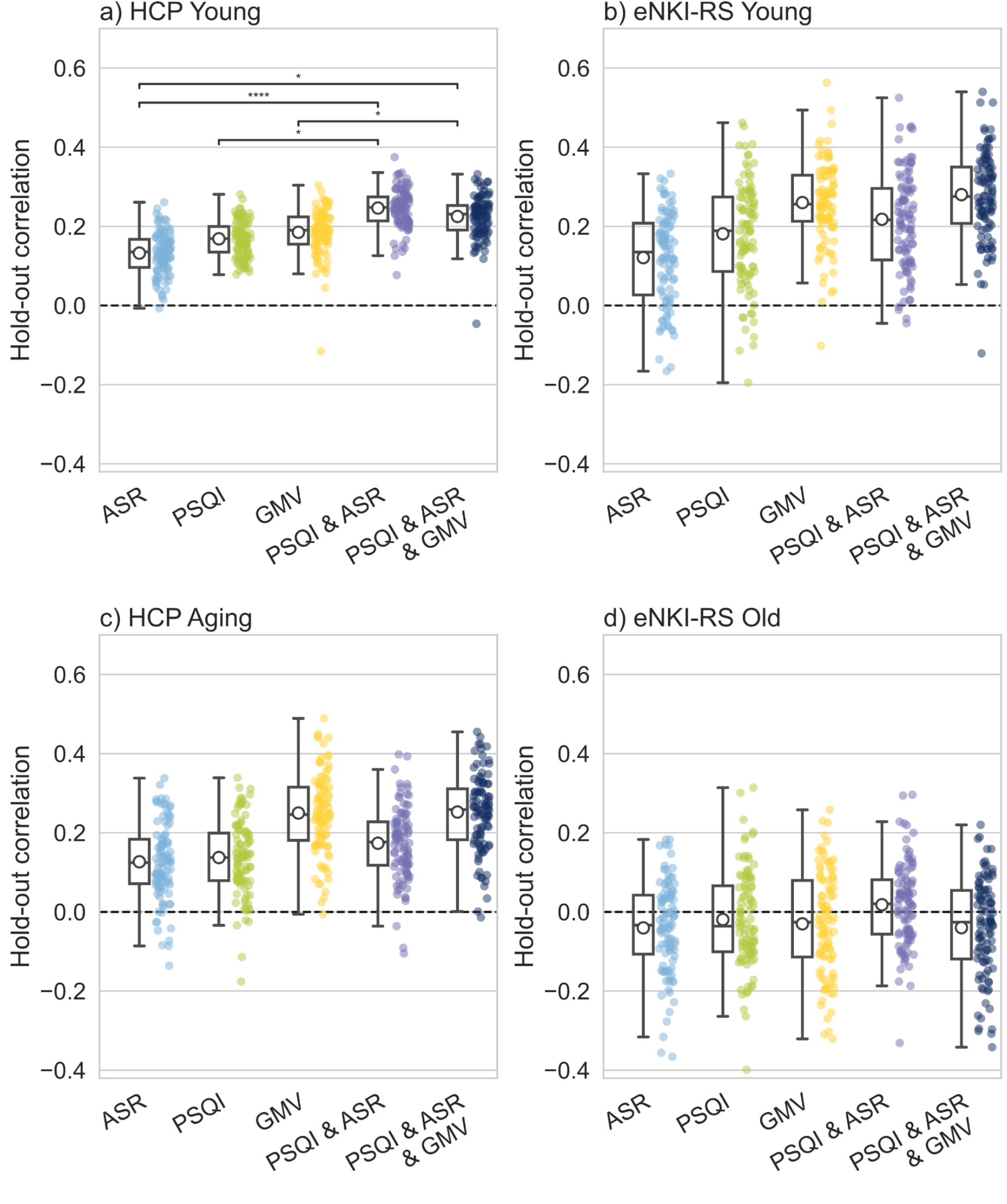
Individual and combined regularized Canonical Correlation Analysis (rCCA) results across four samples. Multivariate associations between depressive symptoms (ASR), sleep quality (PSQI), grey matter volume (GMV), and their combinations with measures of motor performance. Canonical correlation values correspond to the hold-out canonical correlations, computed in 100 outer splits for every model, and in each sample. p-values were obtained from pairwise comparison between models by a corrected resampled t-test, and were corrected for multiple comparisons by False Discovery Rate (FDR) *= p < 0.05, **** = p < 0.0001, missing bars indicate non-significant results.

When sleep quality and depressive symptoms were combined into a single rCCA model, a stronger association with MP was observed compared to the individual models of PSQI and ASR. This enhanced multivariate association reached statistical significance in the HCP Young sample, with a Fisher Z transformed hold-out correlation coefficient of 0.25 ± 0.05. However, in the eNKI-RS Young and HCP Aging samples, we did not observe a significant increase in the correlation coefficients, which remained at 0.22 ± 0.13 and 0.18 ± 0.1, respectively. The model combining PSQI, ASR, and GMV resulted in correlations with MP of 0.23 ± 0.06 (HCP Young), 0.29 ± 0.12 (eNKI-RS Young), 0.26 ± 0.11 (HCP Aging). Therefore, the integrated models consistently yielded stronger or comparably strong associations than individual models (Figure 2a-d).

In our complementary analysis to uncover potential age-independent effects, we combined HCP Young and HCP Aging samples. The model combining PSQI, ASR, and GMV yielded statistically significant higher hold-out correlations compared to all individual models, with a Fisher Z transformed correlation coefficient of 0.22 ± 0.05 (Supplementary Figure 2a). For the redundancy index of Y of the four samples, the integrated models also yielded higher scores than the individual models or were equally high (Supplementary Figure 1). Therefore, we initially present the loadings of the combined model across all domains (Figure 3 and Figure 4). Subsequently, we provided the loadings of the models of GMV and combined PSQI and ASR (Figure 5).

**Figure 3.**
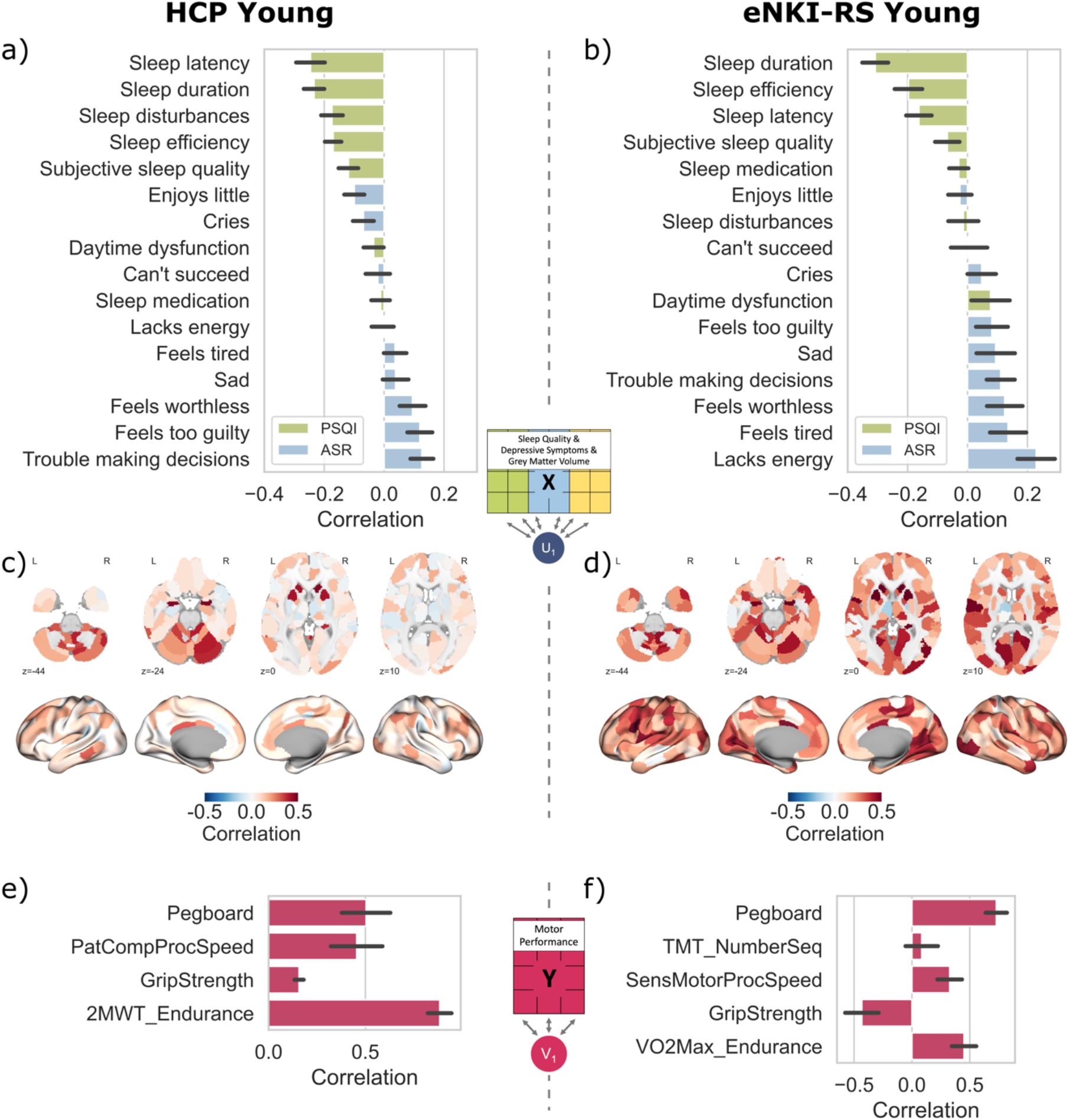
Loadings of regularized Canonical Correlation Analysis of the model combining PSQI, ASR, and GMV vs. motor performance in two samples of young adult. a, b, c, d) All variables of sleep quality (PSQI – Pittsburgh Sleep Quality Index), depressive symptoms (ASR – Adult Self Report), grey-matter-volume parcels are correlated with the canonical variate U; Negative loadings of PSQI and ASR indicate better sleep quality, and less depressive symptoms, while positive loadings indicate worse sleep quality and more depressive symptoms; e, f) All variables of motor performance (Pegboard: 9-hole Pegboard (HCP), Grooved Pegboard (eNKI-RS); PatComp: Pattern Comparison task from NIH toolbox; TMT: Trail Making task; 2MWT: Two-Minute Walk Test; VO2Max: Cardiovascular fitness estimated from bike test) correlated with canonical variate V.

**Figure 4.**
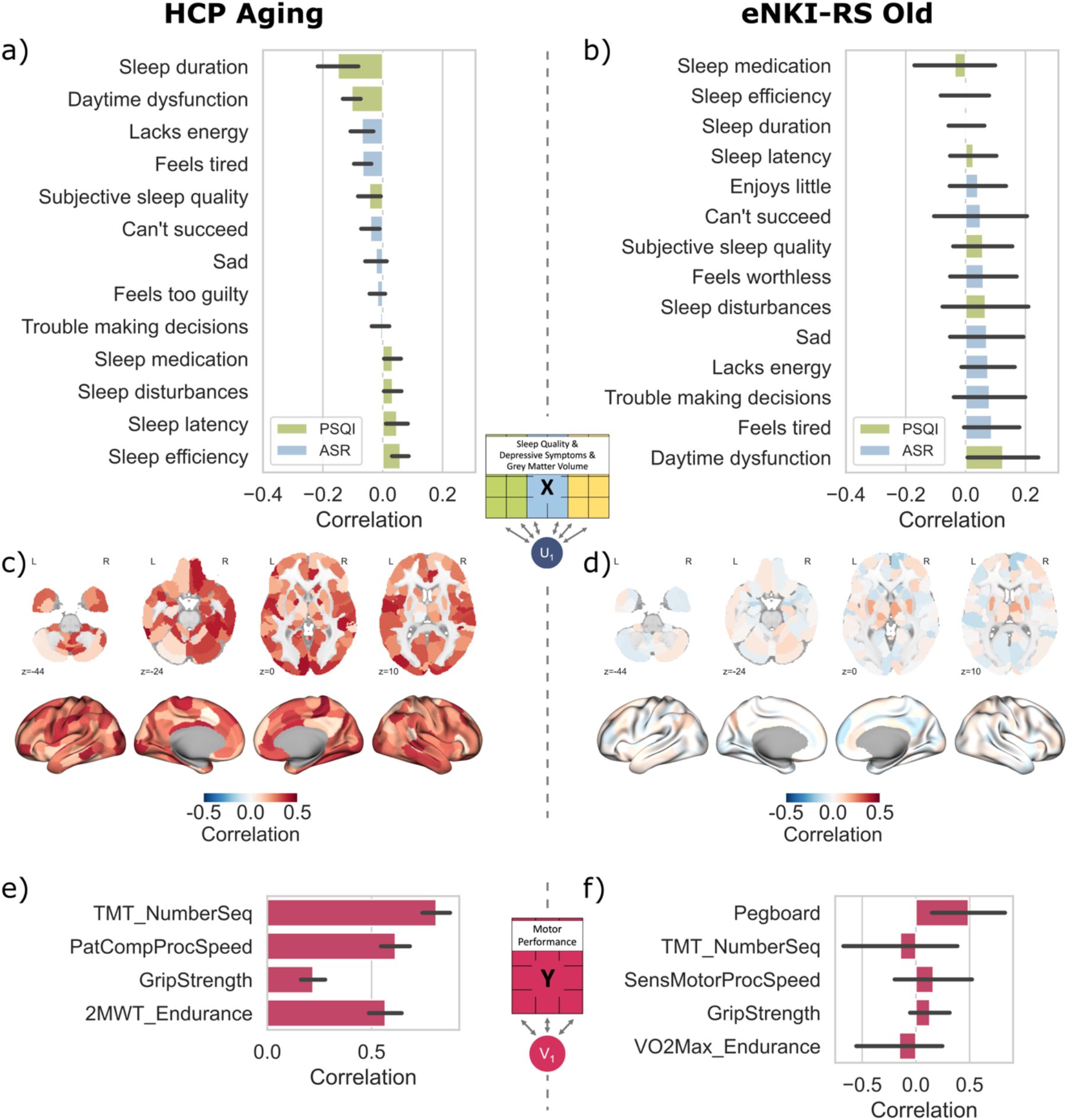
Loadings of regularized Canonical Correlation Analysis of the model combining PSQI, ASR, and GMV vs. motor performance in two samples of older adults. a, b, c, d) All variables of sleep quality (PSQI – Pittsburgh Sleep Quality Index), depressive symptoms (ASR – Adult Self Report), grey-matter-volume parcels are correlated with the canonical variate U; Negative loadings of PSQI and ASR indicate better sleep quality, and less depressive symptoms, while positive loadings indicate worse sleep quality and more depressive symptoms; e, f) All variables of motor performance (Pegboard: Grooved Pegboard; PatComp: Pattern Comparison task from NIH toolbox; TMT: Trail Making task; 2MWT: Two-Minute Walk Test; VO2Max: Cardiovascular fitness estimated from bike test) correlated with canonical variate V. Loadings in the eNKI-RS Old sample show unstable results, averaging around 0.

**Figure 5.**
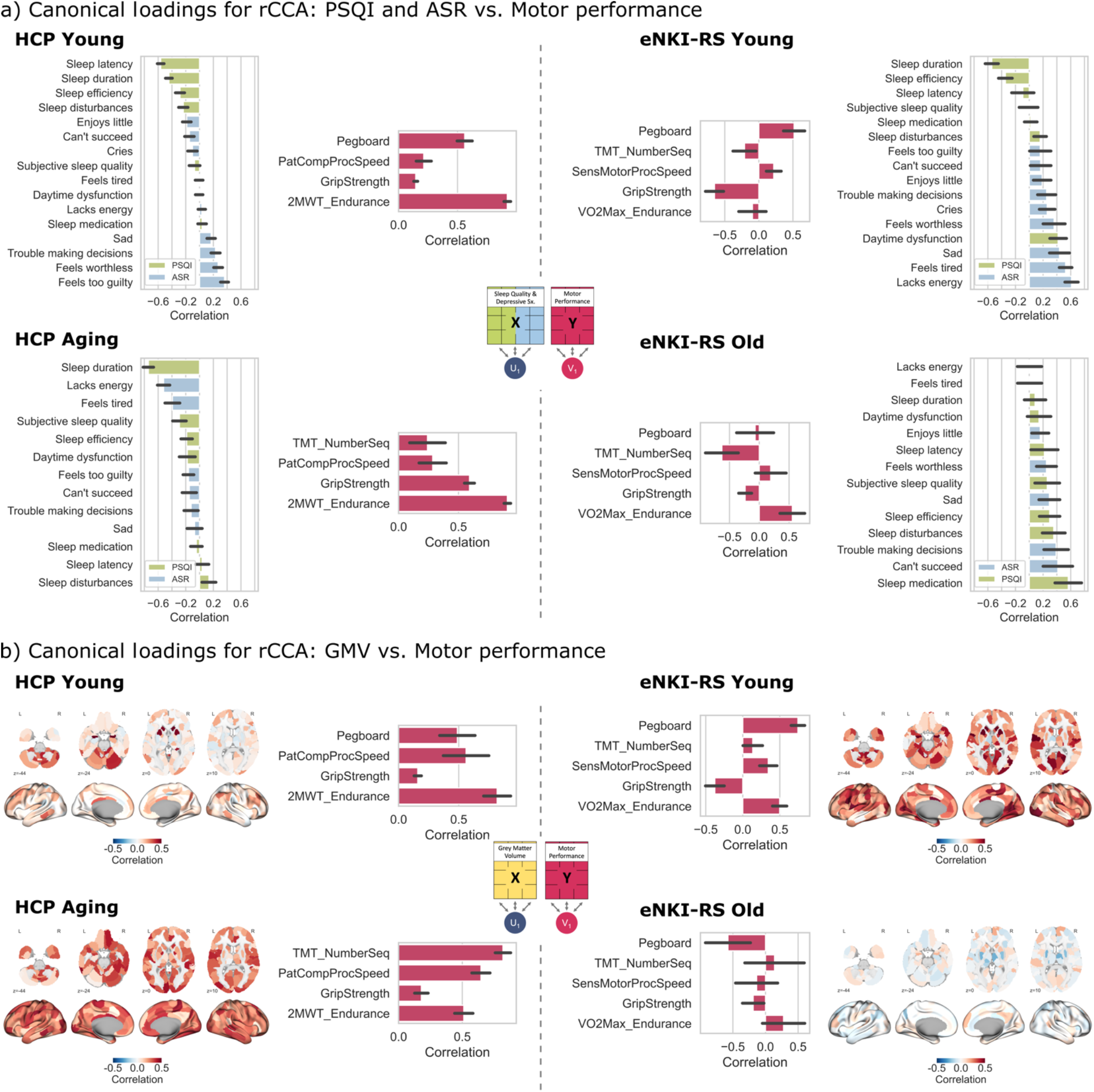
Loadings of regularized Canonical Correlation Analysis of the model combining PSQI & ASR vs. motor performance and GMV vs. motor performance. a) All variables of sleep quality (PSQI – Pittsburgh Sleep Quality Index), depressive symptoms (ASR – Adult Self Report) are correlated with the canonical variate U. Negative loadings of PSQI and ASR indicate better sleep quality, and less depressive symptoms, while positive loadings indicate worse sleep quality and more depressive symptoms; b) All variables of grey-matter-volume parcels are correlated with the canonical variate U. a, b) all variables of motor performance (PatComp: Pattern Comparison task from NIH toolbox; 2MWT: Two-Minute Walk Test; Grip Strength; Grooved Pegboard (eNKI-RS); VO2Max: Cardiovascular fitness estimated from bike test) correlated with canonical variate V.

### Canonical loadings in young adults

To explain the canonical variates of the combined model (PSQI, ASR, GMV vs. MP), we calculated the loadings as Pearson correlations between the input variables and the canonical variates and averaged them over all 100 repeats (Figure 3). The motor canonical variate in the HCP Young sample correlated with endurance (loading = 0.88; measured by the two-minute walk test), dexterity (loading = 0.5; measured by 9-hole Pegboard test), and processing speed (loading = 0.46; measured by Pattern Comparison Task). Grip strength showed the lowest association with the motor canonical variate (loading = 0.16) (Figure 3e). The motor canonical variate in the eNKI-RS Young was also positively associated with dexterity (loading = 0.73; measured by Grooved Pegboard test), processing speed (loading = 0.33; sensory-motor task from Penn Cognitive Task Battery) and endurance (loading = 0.45; by VO2max). Interestingly, grip strength showed a negative association (loading = -0.43) to the motor canonical variate (Figure 3f). This component of higher MP was associated with better sleep quality, coupled with mild depressive symptoms. The loadings of sleep quality and depressive symptoms on the canonical variate showed a consistent pattern between both samples (Figure 3a,b). This was also evident in the significant correlations of the PSQI and ASR loadings (r = 0.83, p<0.01) between the samples. Better sleep quality, particularly self-reported sleep, and bedtime (sleep efficiency: loading = -0.17 (HCP Young), -0.2 (eNKI-RS Young); sleep duration: loading = -0.23 (HCP Young), -0.31 (eNKI-RS Young)), and mild depressive symptoms related to feelings of guilt/self-deprecation (trouble making decisions: loading = 0.13 (HCP Young), 0.11 (eNKI-RS Young)) indicated relevant loadings in both cohorts. In addition, the canonical variate was associated with sleep disturbances in the HCP Young sample and somatic symptoms, lack of energy, and fatigue in the eNKI-RS Young sample. On the brain structural level, we found that various brain regions were associated with MP (Figure 3c,d). The GMV loadings between the two young samples significantly correlated (r_s_=0.31, p<0.0001) with each other. Consistent GMV loadings between both samples were observed in bilateral nucleus accumbens, anterior putamen, lateral amygdala, right mid-cingulate cortex, and parts of the cerebellum. The eNKI-RS Young sample revealed generally higher GMV loadings in a wide range of regions, including the lateral dorsal prefrontal cortex, and inferior parietal lobule.

### Canonical loadings in older adults

The canonical loadings in the combined model (PSQI, ASR, GMV) showed a positive association between sleep quality, depressive symptoms, grey matter volume, and MP in older adults. This association was observed in the HCP Aging sample, but not in the eNKI-RS Old sample (Figure 2d and Figure 4). The motor canonical variate in older adults was composed of the Trail Making Task – Number sequencing (loading = 0.81), Processing Speed Task (loading = 0.61), and the Two-Minute Walk Test (loading = 0.57). Higher MP was associated with better sleep quality, with sleep duration revealing the highest loading (loading = -0.15). Self-reported depressive symptoms, specifically somatic symptoms correlated with better MP (Figure 4a). However, the loadings were relatively low (loading “lacks energy” = 0.07). Relatively high positive loadings of GMV on the canonical variate were observed throughout the brain, suggesting a widespread association of GMV with MP in older adults. Brain regions showing the highest loadings include parts of the cerebellum, left primary motor cortex, right cingulate cortex, bilateral pallidum, and bilateral superior parietal lobule (Figure 4c).

### Canonical loadings of behaviour and brain rCCA

To further disentangle the motor canonical variates associated with behavioural factors (sleep quality, and depressive symptoms) or brain structure (GMV), we analysed the loadings from the respective rCCA models. Notably, the eNKI-RS Old sample did not show robust associations in either analysis (Figure 2d). Thus, while we report the loadings for completeness, they are not discussed further. Statistical comparisons of the loadings between samples showed significant correlations only for GMV. Significant associations were identified between HCP Young and eNKI-RS Young (r_s_=0.27, p<0.0001), HCP Young and HCP Aging (r_s_=0.28, p<0.0001), and eNKI-RS Young and HCP Aging (r_s_=0.39, p<0.0001). Conversely, none of the loadings from the behavioural rCCA models showed significant correlations across any samples.

In a visual comparison of the motor canonical variates of the behavioural rCCA, the highest loadings were observed for the endurance task in the HCP Young and Aging samples, with values of 0.92 and 0.91, respectively (Figure 5a). In the eNKI-RS Young sample, VO2max as a measure of endurance showed no association with the motor component. The pegboard task loaded on the motor component for both HCP Young and eNKI-RS Young, loading = 0.56 and 0.51, respectively. The pegboard task was not measured in the HCP Aging sample. Interestingly, grip strength was associated with the motor canonical variates for both eNKI-RS Young and HCP Aging. In the eNKI-RS Young sample, lower grip strength (r = -0.65) was associated with higher depressive symptoms, especially somatic aspects (Lacks energy r = 0.61, Feels tired r = 0.53). For the HCP Aging sample, higher grip strength (r = 0.59) was linked to longer sleep duration (r = 0.74) and less somatic symptoms (Lacks energy r = -0.52, Feels tired r = -0.39). Processing speed was not really associated with the motor canonical variate in any of the samples. Comparing the motor canonical variates of the brain rCCA models, processing speed showed higher loadings in all three samples (HCP Young = 0.56, eNKI-RS Young = 0.34, HCP Aging = 0.65) (Figure 5b). For the HCP Aging sample, TMT number sequencing was associated with the motor canonical variate (r = 0.82), whereas for eNKI-RS Young this task showed no association. The pegboard task also showed relatively high loadings, with 0.49 in the HCP Young and 0.75 in the eNKI-RS Young samples. Further endurance was associated with the motor component in HCP Young r = 0.82, HCP Aging r = 0.51 and for VO2max in eNKI-RS Young r = 0.5.

## Discussion

The relationship between motor performance (MP) and both behavioural and brain structural factors is complex. Previous studies identified individual associations between those factors, but the multivariate association of sleep quality, depressive symptoms, and GMV with MP has not been extensively investigated. By analysing data from three imaging cohort studies, we found robust and replicable associations between them, particularly in young adults. Higher MP was associated with better sleep quality across the samples. Depressive symptoms showed a differential pattern: young adults showed increased symptoms with higher MP, whereas older adults showed fewer depressive symptoms with higher MP. GMV showed the strongest and most consistent association with MP across all age groups and across individual domains. The combination of all domains into a single rCCA model strengthened the associations. However, rCCA models with sleep quality combined with depressive symptoms or just GMV showed similarly high associations in the HCP Young, and eNKI-RS Young, HCP Aging samples respectively. This suggests two distinct associations of behavioural and brain structural factors with MP.

Shortcomings of this study include the relatively small sample size for brain-behaviour associations (Helmer et al., 2020; Marek et al., 2022). We addressed this issue by using different cohorts but note that different available MP tasks in each cohort only allowed for a conceptual comparison, i.e., a qualitative replication (Xu et al., 2024). The relatively small sample size in one sample of older adults (eNKI-RS Old) may explain the failure to find replicable associations among older adults, but also warrants cautious generalisations. In addition, our analysis method allows the identification of multivariate associations, but in the context of the cross-sectional design, directionality cannot be inferred. Thus, large-scale longitudinal samples across lifespan are required to test the causality between the domains.

### From simple to complex associations of motor performance

Across all models, we found that rCCA models based solely on depressive symptoms had the weakest association with MP (Figure 2). However, models of sleep quality showed a stronger link with MP, and those based on GMV showed the strongest correlations. Interestingly, combining sleep and depressive symptoms improved the canonical correlations and the redundancy index. This differential pattern was significant in the HCP Young and the combined HCP cohorts, suggesting a synergistic pattern. The combined model, including behavioural factors (sleep quality and depressive symptoms) and GMV, significantly increased canonical correlations from the model based on GMV only, but not from the model based on both behavioural domains in the HCP Young sample. However, in the eNKI-RS Young and HCP Aging samples canonical correlations were nearly identical between the combined and GMV only model. Therefore, the association of behaviour and GMV with MP seems dependent on the specific sample, driven either by behavioural factors or brain structure, rather than an additional interaction of the two. This potentially indicates differential modes of association with MP. A possible explanation for these findings is the lack of a common link between brain structure and sleep quality/depressive symptoms, as highlighted previously (Olfati et al., 2024; Weihs et al., 2023; Winter et al., 2024). Similarly, literature assessing the combined effects of brain and behavioural information found no improvement, or even a decrease, in predictability when compared to using only phenotypic information (Dadi et al., 2021; Krämer et al., 2023; Olfati et al., 2024; Omidvarnia et al., 2023). In our study, this is consistent with the results of the HCP Young sample, where the behavioural rCCA model showed the strongest associations to MP and the addition of GMV did not increase the canonical correlation. In contrast, the eNKI-RS Young and HCP Aging samples demonstrated higher hold-out associations and redundancy indices in the rCCA models containing only GMV. This may be due to the influence of other motor tasks or age-related differences.

### Differential brain and behaviour association with MP

We found that a motor canonical variate characterised by endurance, grip strength and dexterity was associated with behavioural factors (Figure 5a). As the motor component correlated very weakly with the more purely cognitive tasks (processing speed and TMT number sequencing), it may represent a more fundamental component of motor function. Endurance showed the highest loadings in both HCP cohorts, but not in the eNKI-RS Young sample, possibly due to various motor tasks. Moreover, short sleep duration was associated with worse MP, which aligns with literature linking sleep deprivation and poor sleep quality with lower endurance (Craven et al., 2022). Cardiovascular endurance was further linked to sleep latency in a cohort of middle-aged and older adults, which we similarly observed in the HCP Young sample (Hsu et al., 2021). Grip strength showed high loadings in the eNKI-RS Young and HCP Aging samples, positively associated with fewer depressive symptoms. This is in line with previous findings (Gu et al., 2021). Interestingly, a recent study found that short sleep duration increased the risk for developing depressive symptoms, but this effect was attenuated in participants with high grip strength (R. Chen et al., 2023). A study of a mixed cohort of depressed and non-depressed people found that the pegboard task and other measures of processing speed were positively related to depressive symptom severity, but not the diagnosis of depression (Shura et al., 2017). In contrast to these results and our expectations, we observed mild depressive symptoms being associated with higher MP in young adults. Possible reasons for this remain speculative but could include self-doubt as a potential motivational factor (Ede et al., 2017), higher anxiety due to sleep disturbance or perfectionism (Riemann et al., 2020), or increased exercise as a compensatory strategy in those with generally higher depressive symptoms (Noetel et al., 2024), given that the participants were from the general population and did not meet diagnostic criteria for clinical depression. Conversely, in the HCP Aging sample, we observed low depressive symptoms with higher endurance and grip strength, which is consistent with previous literature (Gu et al., 2021; Kandola et al., 2020; Kitagaki et al., 2020). The results from our multivariate approach showed that a combination of normal sleep duration and efficiency, together with fewer somatic symptoms, as well as mild affective aspects, are associated with higher MP.

The association with GMV showed a motor canonical variate driven more by cognitive aspects of processing speed, dexterity, and endurance (Figure 5b). Previous studies have identified associations between MP and GMV in a wide range of regions. Macroanatomical brain correlates with manual MP were found for cerebral GMV, but not consistently for cerebellar GMV (Hoogendam et al., 2014; Koppelmans et al., 2015). A recent study found associations between GMV atrophy and manual dexterity in more fine-grained parietal areas and gross motor function in the temporal regions (Dougherty et al., 2024). Fitness and physical activity have been mostly associated with GMV differences in the prefrontal cortex and the hippocampus (Erickson et al., 2014). In this study, we found high loadings in a number of cortical regions in the combined rCCA of young adults, such as the right mid-cingulate cortex, lateral dorsal prefrontal cortex, inferior parietal lobule. The combined rCCA of older adults showed loadings in the left primary motor cortex, right cingulate cortex, bilateral superior parietal lobule. These regions were associated with better MP. Furthermore, rCCA loadings in subcortical regions, including bilateral nucleus accumbens, anterior putamen, lateral amygdala for young adults, and bilateral pallidum for older adults, and multiple cerebellar regions were observed.

### Different associations with MP in young and old adults

The relationship between the behavioural and brain domains with MP exhibited distinct patterns between younger and older adults (Figure 3 and Figure 4). In younger adults, better sleep characteristics, such as increased duration, reduced latency, and higher sleep efficiency together with mild depressive symptoms, were associated with higher MP. In contrast, older adults demonstrated that primarily sleep duration below 7 hours and less somatic-related depressive symptoms correlated with better MP.

Age-related increases in between-participant variability could potentially explain the failure of the second sample of older adults to generalise (Hunter et al., 2016). Ageing affects MP through multiple aspects, such as sarcopenia, mobility issues, cardiovascular and metabolic health, and cognitive decline (Jin et al., 2023; Tai et al., 2022; Tieland et al., 2018). Moreover, ageing differently impacts sleep in older adults, the sleep becomes more fragmented, sleep latency increases, and slow-wave sleep decreases (Mander et al., 2017). In addition, the nature of depressive symptoms shifts in older people, with an increased prevalence of somatic complaints, whereas feelings of guilt may be more prevalent in younger adults (Abrams & Mehta, 2019; Hegeman et al., 2012). These age-related differences highlight the complex interplay of biological and behavioural factors affecting MP across the lifespan.

## Conclusion

Our findings revealed a distinct pattern of associations between behavioural and brain structural factors with MP. Using a machine learning framework to ensure robustness of our results, we found that better sleep quality and mild depressive symptoms were associated with better MP in young adults. This was conceptually replicated in a second young cohort. In a cohort of older adults, we observed that healthy sleep and fewer depressive symptoms were associated with better MP. Brain-related associations highlighted more cognitive aspects of MP. We hope that these findings increase incentive regarding the importance of sleep quality and depressive symptoms to improve motor functioning of ordinary individuals in the society, professional athletes, patients with motor-related neurodegenerative diseases, such as Parkinson’s disease, as well as patients with psychiatric conditions including major depressive disorder. In order to extend these findings and assess the reproducibility of our findings, future large-scale studies using open data sharing (Ganz & Poldrack, 2023) and ENIGMA consortium (Schmaal et al., 2020; Tahmasian et al., 2021; Thompson et al., 2020) should assess a range of motor behaviours together with a broader range of phenotypic measures, such as personality, environmental and lifestyle factors in younger and older adults, which are critical aspects toward personalised medicine (Delpierre & Lefèvre, 2023). In particular, longitudinal assessments would help in elucidating the causal relationships between behavioural and brain structural aspects of MP.

## Acknowledgment

VK is supported by the Deutsche Forschungsgemeinschaft (DFG, German Research Foundation) – Project-ID 431549029 – SFB 1451. SBE received Helmholtz Imaging Platform grant (NimRLS, ZT-I-PF-4-010). BTTY is supported by the NUS Yong Loo Lin School of Medicine (NUHSRO/2020/124/TMR/LOA), the Singapore National Medical Research Council (NMRC) LCG (OFLCG19May-0035), NMRC CTG-IIT (CTGIIT23jan-0001), NMRC STaR (STaR20nov-0003), Singapore Ministry of Health (MOH) Centre Grant (CG21APR1009), the Temasek Foundation (TF2223-IMH-01), and the United States National Institutes of Health (R01MH120080 & R01MH133334).

Data were provided [in part] by the Human Connectome Project, WU-Minn Consortium (Principal Investigators: David Van Essen and Kamil Ugurbil; 1U54MH091657) funded by the 16 NIH Institutes and Centers that support the NIH Blueprint for Neuroscience Research; and by the McDonnell Center for Systems Neuroscience at Washington University. Research reported in this publication was supported by the National Institute On Aging of the National Institutes of Health under Award Number U01AG052564 and by funds provided by the McDonnell Center for Systems Neuroscience at Washington University in St. Louis. The HCP-Aging 2.0 Release data used in this report came from DOI: 10.15154/1520707.

Data used in the preparation of this manuscript were obtained from the National Institute of Mental Health (NIMH) Data Archive (NDA). NDA is a collaborative informatics system created by the National Institutes of Health to provide a national resource to support and accelerate research in mental health. Dataset identifier(s): NIMH Data Archive Digital Object Identifier (DOI) 10.15154/qq04-sd33. This manuscript reflects the views of the authors and may not reflect the opinions or views of the NIH or of the Submitters submitting original data to NDA.

## Author Contributions

Conceptualization: VK, AD, SBE, MT; Software: VK, HB, FH; Formal analysis: VK, FH; Data Curation: VK, FH; Writing – Original Draft: VK, MT; Writing – Review & Editing: VK, HB, ENS, FH, BTTY, AD, SBE, MT; Visualization: VK; Supervision: AD, SBE, MT

## Data and Code Availability

We used publicly available datasets. Access to the Human Connectome Project can be requested after registering and accepting the data-use terms (db.humanconnectome.org). Access to the HCP Aging can be requested and accessed through the NIMH Data Archive, further information can be found on website of the HCP (https://www.humanconnectome.org/study/hcp-lifespan-aging/data-releases). Data of the enhanced Nathan Kline Institute Rockland Sample can be accessed as outlined here: https://fcon_1000.projects.nitrc.org/indi/enhanced/access.html.

The code used for the rCCA and machine learning framework computation is available at https://github.com/anaston/cca_pls_toolkit. CAT12 can be downloaded at https://neuro-jena.github.io/cat/index.html#DOWNLOAD.

