## Supplemental Material for "Lower motor performance is linked with poor sleep quality, depressive symptoms, and grey matter volume alterations"

### Supplemental Methods

#### Participants and phenotypic data

Data from three different publicly available cohorts was analysed: the Human Connectome Project Young Adult (HCP Young, S1200 release), the Human Connectome Project Aging (HCP Aging, 2.0 release), and the enhanced Nathan Kline Institute-Rockland Sample (eNKI-RS) (Bookheimer et al., 2019; Nooner et al., 2012; Van Essen et al., 2013). To investigate potential age-related differences, participants were divided into younger (HCP Young: 22-37 years; eNKI-RS Young: 18-40 years) and older adults (HCP Aging: 50-85 years, eNKI-RS Old: 50-85 years). After excluding participants with missing data and low-quality neuroimaging data, a total of 1950 participants (1112 women) were included.

**HCP Young**. 1086 participants (587 female, 22-37 years). Sleep quality was assessed using 7 components computed in the Pittsburgh Sleep Quality Index (PSQI) (Buysse et al., 1989) (Buysse et al., 1989). Depressive symptoms were measured using the relevant items of the Adult Self-Report (ASR) for ages 18-59 associated with depressive Disorder (Achenbach, 2013; Achenbach & Rescorla, 2003). Items not included in the ASR version for older adults for ages 60+ (OASR) and sleep-related questions were excluded. Motor-related phenotypes were acquired using NIH Toolbox normalised scores for (1) Grip strength (Strength), an assessment of upper body strength using a dynamometer, (2) 2 Minute Walk Test (Endurance), a test of physical fitness, cardiovascular endurance by measuring the distance covered by a participant in 2 minutes time, (3) 9-Hole Pegboard Test (Dexterity), an assessment of fine motor dexterity by measuring the time that it takes participants to place pegs in 9 holes and remove them (Reuben et al., 2013), and (4) Pattern Completion Processing Speed (Processing Speed), an assessment of the speed of processing by asking participants to identify as quickly as possible if two pictures are the same or not (Weintraub et al., 2013). For ASR and PSQI higher scores indicate higher symptom severity, i.e., more depressive symptoms and worse sleep quality. For all motor measures higher scores indicate better motor performance, i.e., stronger, faster.

**eNKI-RS Young**. 230 participants (128 female, 18-40 years). Sleep quality and depressive symptoms were measured as in HCP Young. To make the MP measures comparable to the human connectome project, we took the (1) grip strength of the dominant hand (Strength), (2) VO2max by bike test (Endurance), a more direct measurement of cardiorespiratory fitness by measuring the heart rate on a stationary bike (3) Grooved Pegboard Test (Dexterity), similar to the 9-hole pegboard but the participants are required to place 25 pegs, which are uniquely shaped (4) Mouse Practice task (Processing Speed), a test in which participants have to react as quickly as possible to green squares appearing on a computer screen by moving the cursor and clicking on them (5) Trail Making Test A (TMT Number Seq – Sensorimotor Speed), participants are asked to connect 25 numbered dots (1,2…,25) as quickly as possible (Åstrand & Ryhming, 1954; Delis et al., 2001; Gur, 2001; Reuben et al., 2013). It should be noted that the VO2max calculated by the bike test could be considered as an indirect measure of cardiorespiratory fitness as well. The submaximal VO2max and finger tapping tasks were both excluded due to the relatively small number of participants who performed these measures. In addition, TMT Number Seq was chosen instead of TMT Motor speed of the Delis-Kaplan to obtain a measure comparable to HCP Aging TMT Number Seq.

**HCP Aging**. 354 participants (198 female, 50-85 years). Sleep quality was measured by the PSQI questionnaire, and depressive symptoms by the OASR as described above. MP measures are based on NIH Toolbox normalised scores for Grip strength (Strength), Minute Walk Test (Endurance), Pattern Completion Processing Speed (Processing Speed), and raw score of Trail Making Test A (Reitan, 1992; Reuben et al., 2013; Weintraub et al., 2013).

**eNKI-RS Old**. 280 participants (199 female, 50-85 years). Sleep quality and depressive symptoms were measured as in HCP Aging. Motor related phenotypes were identical to those of the eNKI-RS Young sample.

#### Supplemental Results

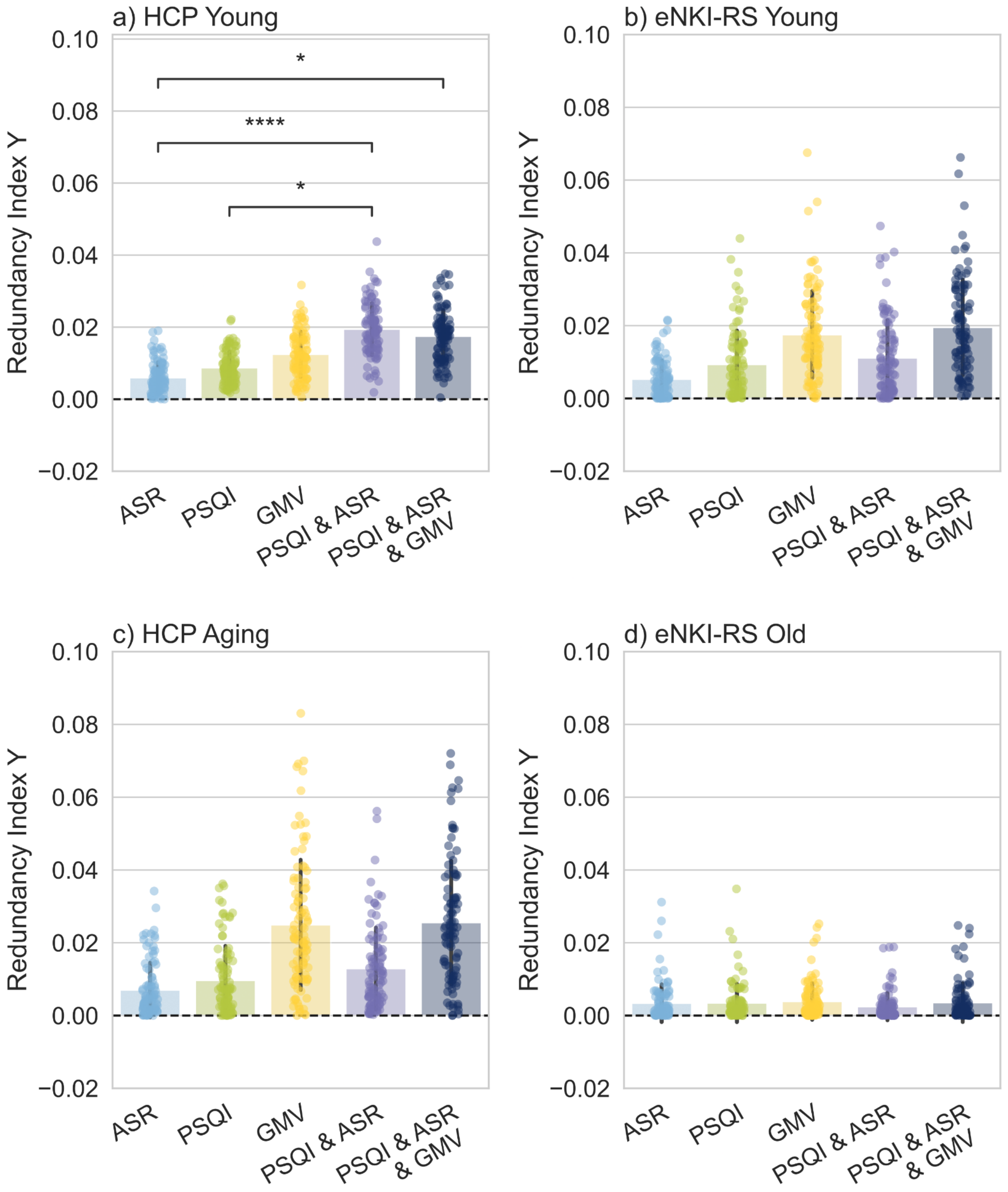


***Supplementary Figure 1.*** *Redundancy Index of Y. Model comparison based on machine learning adapted t-test. p-values are FDR corrected.*

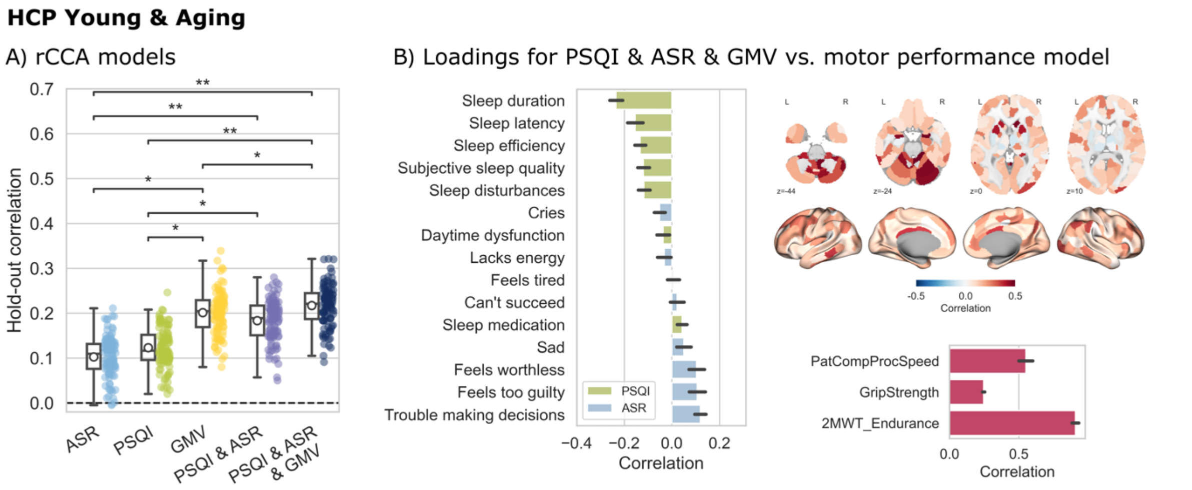


***Supplementary Figure 2.*** *Combined sample of HCP Young and Aging cohort (n = 1631, 894 female, age range = 22-85). A) Individual and combined regularized Canonical Correlation Analysis (rCCA) 100 times repeated hold-out correlations. Pairwise comparison between models within the sample by machine-learning adjusted t-test, p-values are False Discovery Rate (FDR) corrected, *= p < 0.05, ** = p < 0.001, missing bars indicate non-significant results. B) Canonical loadings of rCCA of the model combining PSQI & ASR & GMV vs. motor performance. All variables of sleep quality (PSQI – Pittsburgh Sleep Quality Index), depressive symptoms (ASR – Adult Self Report), grey-matter-volume parcels are correlated with the canonical variate U, negative loadings of PSQI and ASR indicate better sleep quality, and less depressive symptoms, while positive loadings indicate worse sleep quality and more depressive symptoms; and all variables of motor performance (PatComp: Pattern Comparison task from NIH toolbox; 2MWT: Two-Minute Walk Test; Grip Strength) correlated with canonical variate V.*
